# Informing biologically relevant signal from spatial transcriptomic data

**DOI:** 10.1101/2024.09.09.610361

**Authors:** Emir Radkevich, Darwin D’Souza, Raphael Mattiuz, Raphael Merand, Rachel Chen, Jarod Morgenroth-Rebin, Krista Angeliadis, Daniella Nelson, Anis Kara, Travis Dawson, Igor De Souza, Kai Nie, Pauline Hamon, Samarth Hegde, Jesse Boumelha, Rachel Brody, Sinem Ozbey, Clotilde Hennequin, Di Feng, Jie Dai, Edgar Gonzalez-Kozlova, Zhihong Chen, Seunghee Kim-Schulze, Sacha Gnjatic, Miriam Merad, Vladimir Roudko

**Affiliations:** Department of Immunology and Immunotherapy, Icahn School of Medicine at Mount Sinai, New York, NY, 10029, USA; Human Immune Monitoring Center, Icahn School of Medicine at Mount Sinai, New York, NY, 10029, USA; Department of Pathology, Icahn School of Medicine at Mount Sinai, New York, NY, 10029, USA; Department of Oncological Sciences, Icahn School of Medicine at Mount Sinai, New York, NY, 10029, USA; Mount Sinai Biorepository, Department of Pathology, Icahn School of Medicine at Mount Sinai, New York, NY, 10029, USA; The Marc and Jennifer Lipschultz Precision Immunology Institute, Icahn School of Medicine at Mount Sinai, New York, NY, 10029, USA; Department of Oncological Sciences, Tisch Cancer Institute, Icahn School of Medicine at Mount Sinai, New York, NY, 10029, USA; Global Computational Biology and Digital Sciences, Boehringer Ingelheim Pharmaceuticals, Inc., 900 Ridgebury Road, Ridgefield, CT, 06877, USA; Cancer Immunology and Immune Modulation, Boehringer Ingelheim Pharmaceuticals, Inc., 900 Ridgebury Road, Ridgefield, CT, 06877, USA

## Abstract

Visium is a spatial sequencing technology that utilizes messenger RNA (mRNA) to spatially map gene expression within tissues. Despite its potential, research utilizing deconvolution tools and exploring microenvironment dynamics remains challenging. We address this gap by benchmarking deconvolution tools across diverse biological contexts, identifying optimal methodologies. Subsequently, we introduce a novel pipeline integrating advanced deconvolution techniques and novel tools for reproducible tissue microenvironment analysis. Through this approach, we uncover intricate immune aggregate biology, highlighting the power of our methodology in unraveling complex biological phenomena.

## INTRODUCTION

Spatial transcriptomics has emerged as a transformative technology, revolutionizing the way we perceive and analyze gene expression within the context of tissue architecture. At its core, spatial transcriptomics integrates high-throughput sequencing techniques with spatial information, enabling the visualization and quantification of gene expression patterns within intact tissue sections. By preserving the spatial context of gene expression, this approach transcends traditional bulk, offering unprecedented insights into cellular heterogeneity, tissue microenvironments, and spatial organization of gene expression networks.

The impact of spatial transcriptomics on biological research is multifaceted and far-reaching. First, it facilitates the elucidation of cellular interactions and signaling dynamics within tissues, unraveling the intricate interplay between different cell types in health and disease. Second, it provides spatially resolved molecular maps that aid in the identification of novel cell populations, biomarkers, and therapeutic targets, thereby accelerating drug discovery and development processes. Third, spatial transcriptomics enables the characterization of spatially restricted gene expression programs underlying developmental processes, tissue morphogenesis, and regenerative capacities ^1^.

Key methodologies encompassing spatial transcriptomics include in situ sequencing ^2^, spatially resolved transcriptomics technologies such as Slide sequencing [Slide-seq] ^3^, Visium ^4,5^ and Visium HD ^6^, and imaging-based approaches like single-molecule fluorescence in situ hybridization [smFISH] Merfish ^7^, CosMX ^8^, Xenium ^9^.

Slide-seq, harnessing spatially barcoded beads for messenger RNA (mRNA) capture, offers a scalable approach with remarkable sensitivity, facilitating the analysis of transcriptome-wide readout. However, its spatial resolution may not match the precision achieved by other techniques due to bead diffusion which could potentially compromise spatial fidelity. Merfish, a method capable of multiplexing detection of hundreds to thousands of RNA species at single-molecule resolution within intact tissues, stands out for its unprecedented spatial and molecular granularity. Nevertheless, its adoption is hindered by the intricate experimental protocols and specialized instrumentation required. CosMX, integrating multiplexed imaging of RNA and proteins using unique cellular barcodes, promises comprehensive insights into tissue architecture. Yet, the technique’s multiplexing capacity and probe optimization necessitate careful consideration for effective implementation. Xenium, a technique leveraging spatially barcoded DNA nanostructures for mRNA profiling using disease specific probe panels, exhibits promise for high-resolution spatial transcriptomic analysis with minimized background noise. However, its scalability necessitates further scrutiny and validation. Moreover, all listed smFISH methods capture only a relatively short panel of genes compared to other available transcriptome-wide approaches, for example Slide-seq and Visium.

In contrast, Visium has been by far the most adopted technology across academic institutions around the world ^10^. Visium employs spatially barcoded oligonucleotides to capture mRNA from tissue sections, enabling the analysis of thousands of genes with whole transcriptome resolution. This balance between sensitivity, resolution, and accessibility has positioned Visium as a cornerstone in spatial transcriptomic research across diverse scientific domains, offering profound insights into the spatial orchestration of gene expression within complex biological landscapes ^11^.

Nevertheless, Visium has a few limitations such as limited spatial coverage compared to Xenium, relatively low spatial resolution (the lowest bit of information is spot of 55 micrometer diameter) and lower per-gene sensitivity compared to Xenium. At the time of writing this paper, 10X Genomics introduced Visium HD technology, which offers single-cell resolution with an 8-micrometer bin. However, there is currently only one available technical preprint comparing Visium and Visium HD ^6^. Therefore, further research is necessary to validate and understand the full implications of this new technology.

There is also a gap in available computational tools describing the microenvironment of a selected tissue region in Visium samples. While there are tools such as squidpy Python package ^12^, that identify regions that are often close to each other, share similar architecture or calculate co-occurrence of tissue regions, there is no tool developed to our knowledge that can describe the microenvironment of a selected tissue region.

Our algorithm identifies Visium spots positioned equidistantly from a selected structural boundary, whether they are inside the structure (internal layers) or moving away from it (outer layers). Then, we regress gene expression or cellular counts across these layers and thus can explore the microenvironment of a selected tissue region. Although spatial resolution of Visium is not at the single cell level, we showcase how spot deconvolution along with our developed tool can help break down the complexity of the tumor tissue microenvironment.

## RESULTS

### Deconvolution of Visium spots is an essential step in Visium downstream analysis

Spatial transcriptomic datasets possess a unique challenge both on acquisition and analysis sides of the workflow. To address these unique challenges, we developed an analytical pipeline aiming to reliably detect and quantify spatially resolved biomarkers (**Figure 1A**). The approach combines below research strategies:

1. Annotation by pathologist to get ground-truth of tissue areas matched across the dataset samples;
2. Hematoxylin and eosin (H&E) based cell segmentation to quantify number of cells per each Visium spot using StarDist ^13,14^;
3. Optional – perform image-based classification of cell type lineages considering the cellular shape after segmentation as a predictor;
4. Quality control (QC), batch correction and integration of transcriptome dataset;
5. Transcript mixture modeling and projection on tissue-matched annotated single cell dataset (deconvolution) and multiplication on total number of segmented cells to extract absolute quantities of deconvoluted cell types;
6. Pathway analysis of spatially resolved transcriptome data;
7. Statistical and survival analysis using cellular biomarkers: quantities per area; normalized pathway enrichments per area; gradient analysis over the borders between two adjacent areas of interest.

To validate step 5 (deconvolution) we performed benchmarking of available deconvolution tools comparing the results to pathologist-based annotated areas. In general, profiled tools [Cell2location ^15^, Stereoscope ^16^, RCTD ^17^, Seurat ^18^, Tangram ^19^, SpatialDWLS ^20^, DestVI ^21^] show similar true positive rate as shown by the ROC curve analysis per pathological area (**Figure 1B, Supplementary Figure S1**) as well as overall correlation of proportions per tissue type (**Supplementary Figures S2)**. Cell2location results were used for all downstream analysis.

**Figure 1.**
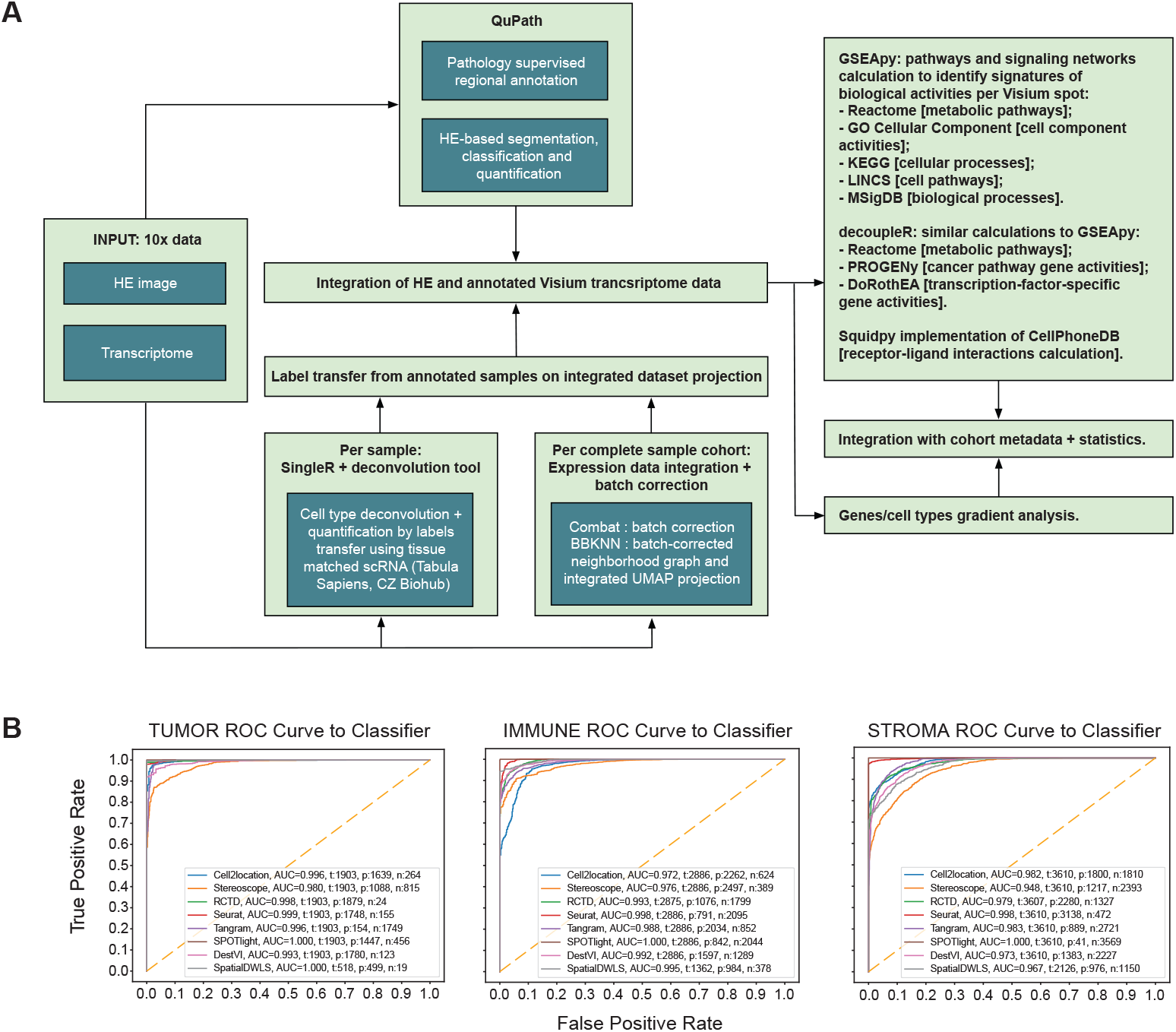
Deconvolution as an essential part of Visium analysis. (A) Schema of the developed Visium pipeline. (B) Receiver Operating Characteristic (ROC) curves are plotted for each tool within a pathological area of colorectal cancer tissue from which Area Under the Curve (AUC) scores were subsequently calculated. Deconvolution outputs were compared against ground truth spot labels expertly annotated by a trained pathologist. Ground truth labels were classified as either Tumor, Immune or Stroma. Legend of the ROC curves show for each tool the AUC score, total spots (t), positive spots (p), meaning labels that were concordant between the deconvolution tool and pathologist and vice versa with negative spots (n).

### Clustering using deconvolution matrix yields superior results comparing to gene expression matrix

We applied the pipeline (**Figure 1A**) to colorectal cancer human samples (CRC sample C1, C2 and C3) using recently published single cell colorectal dataset as a source of cell references for deconvolution step ^22^.

We noticed that clustering absolute numbers of cells projected in each Visium spot using Leiden clustering algorithm ^23^ revealed tissue patterns not observed using gene expression matrix as a data source (**Figure 2A**). For example, sample C1 demonstrates the superiority of a deconvolution matrix as a source of input data since an additional immune aggregate (cluster 3 in deconvolution column) was found and subsequently validated with the pathologist as an immune aggregate. In the sample C2 both approaches result in finding an immune aggregate region (clusters 7) and immune-cells-enriched compartment (clusters 1 and 0 in gene-expression-derived and deconvolution-derived respectively). In the sample C3 deconvolution-based approach shows better granularity of tumor and immune compartments (clusters 5/7 and 6 respectively) compared to gene expression approach. Thus, clustering absolute number of cells (deconvolution results) is superior to results obtained from clustering gene counts (**Figure 2A**).

**Figure 2.**
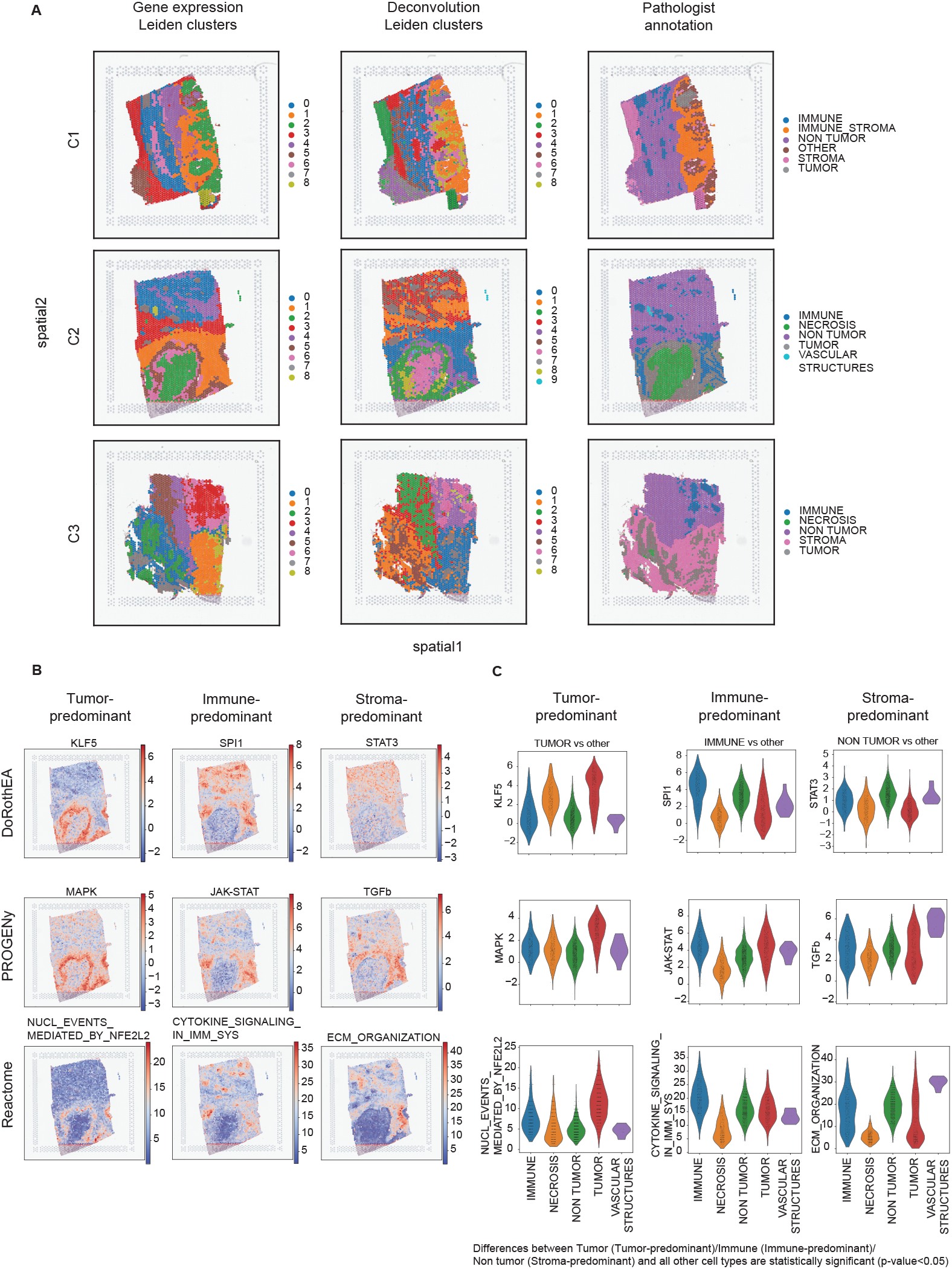
Clustering of deconvolution matrices is more beneficial than clustering gene expression matrices only. (A) Three row panels show the results of unsupervised Leiden clustering performed on gene expression matrix (n=17943 genes; first column) or deconvolution matrix (n=98 cell types; second column) and the results of pathologist annotation. Each row represents one colorectal human sample. Sample C1 demonstrates the superiority of a deconvolution matrix as a source of input data for Leiden clustering since an additional immune aggregate (cluster 3 in deconvolution column) was found and subsequently validated with the pathologist as an immune aggregate. In the sample C2 both approaches result in finding an immune aggregate region (clusters 7) and immune-cells-enriched compartment (clusters 1 and 0 in gene-expression-derived and deconvolution-derived respectively). In the sample C3 deconvolution-based approach shows better granularity of tumor and immune compartments (clusters 5/7 and 6 respectively) compared to gene expression approach. (B) Unsupervised pathway activity analysis identifies pathways that resemble pathologist annotated histological areas. (C) Quantification of pathway activity scores across pathologist annotated histological areas. All the differences between Tumor and other compartments in “Tumor vs other”, Immune and other compartments in “Immune vs other” and Non-tumor and other compartments in “Non-immune vs other” are statistically significant (Mann-Whitney U test, p-value<0.05).

All the subsequent analysis steps were performed on all three colorectal samples and yielded similar results. In this paper we show sample C2 as an example.

We performed functional analysis on Visium data and discovered close resemblance of some pathways to gross histological annotation. We used three different databases – DoRothEA ^24^, PROGENy ^25^ and Reactome ^26^ – as a source of pathways and found nine pathways, that can be categorized into tumor-, immune- and stroma-related, described different regions in the C2 CRC sample (**Figure 2B**). Projecting these pathways and quantifying distribution of their activity scores confirmed selective enrichment of these pathways within pathology-matched areas. STAT3, TGFb and “extracellular matrix organization” signatures are highly predominant in stromal region; JAK-STAT, “cytokine signaling” and SPI1 – in immune region; MAPK, NFE2L2 and KLF5 – in tumor region [Mann-Whitney U-test, p-value<0.05] (**Figure 2C**).

### Deconvolution of Visium spots uncovers intricate biology of tertiary lymphoid structures

Next, we hypothesized whether our approach of unsupervised cell-composition clustering of immune aggregates can reveal and quantify heterogeneity within them as well as parse out differences between spatially isolated aggregates or not. We performed clustering with increased resolution on colorectal sample C2. Indeed, increasing resolution of clustering lead to immune aggregate split on two distinct patterns; “core” (matched with cluster 11) and “border” (matched with cluster 6), each of them having distinct cellular composition: “core” is enriched in GC-type B cells, mregDCs, Tfh cells, while “border” – in IgG and IgA plasma cells. This pattern closely resembled tertiary lymphoid structure (TLS) – immune aggregates with ongoing spatially resolved antigen-specific immune reactions (**Figure 3A**).

**Figure 3.**
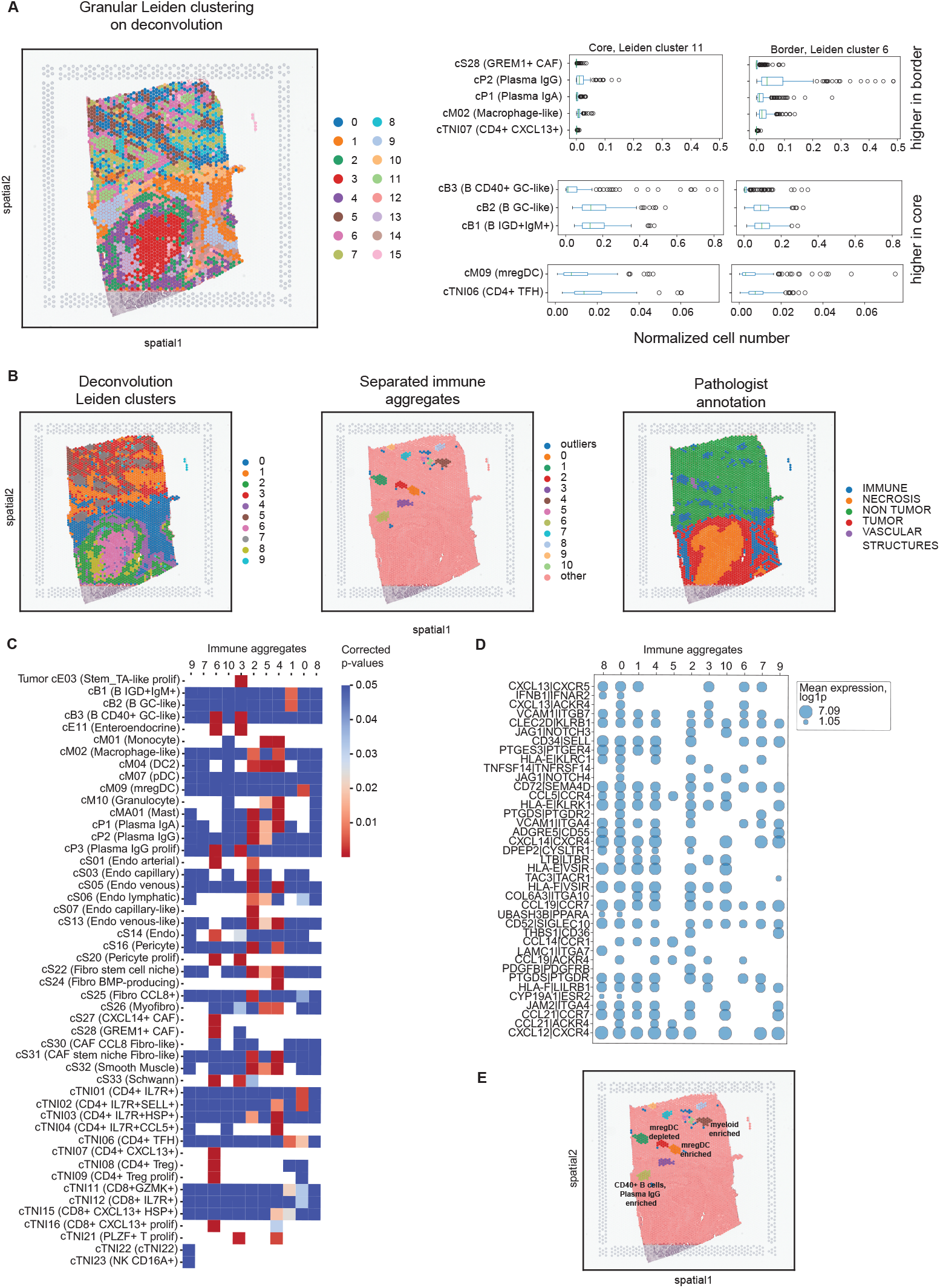
Immune aggregates structural study shows the major differences among aggregates on a histological slide. (A) Left panel shows the results of unsupervised Leiden clustering on a histological slide (see sample C2 on Figure 2), where structures resembling immune aggregates are divided into core and border parts. Right panel highlights cell types driving the difference between core and border compartments (Mann-Whitney U test, p-value<0.05, Benjamini-Hochberg multiple correction is applied). Normalization is performed by the mean number of cells per spot for each area. (B) Density-based spatial clustering (DBSCAN) reveals fine separation of immune aggregates on one histological slide. (C) Heatmap of corrected p-values testing one immune aggregate vs other aggregates (Mann-Whitney U test, ‘greater’ alternative, Benjamini-Hochberg multiple correction). Immune aggregates 2, 4, 3, 5 and 6 differ from the other immune aggregates majorly since these aggregates have the biggest number of cell types driving the difference. (D) Calculated ligand-receptors interactions inside immune aggregates using CellPhoneDB. Aggregates 3, 5, 6 and 7 exhibit different patterns of ligand-receptor pairs. log1p is the natural logarithm of one plus the input array. (E) Labeled immune aggregates highlighting major descriptive features.

To understand TLSs at higher resolution at the cell types and frequencies, we used density-based spatial clustering (DBSCAN) ^27^ which allows to split geographically isolated communities belonging to the same annotation type. Doing so, we were able to isolate 11 TLSs within C2 CRC sample (**Figure 3B**). We performed Mann-Whitney U test “one vs other” (p-value<0.05) on all TLSs and described the cell types driving the difference between the structures (**Figure 3C, Supplementary Figure S3**). For example, we found immune aggregate “0” to be mregDC-enriched while “2” to be mregDC-depleted (**Supplementary Figure S3B**).

Additional analysis of ligand-receptor interaction on separated immune aggregates gave us more information about the nature of underlying processes in TLSs (**Figure 3D**). For example, cytokines of CXCL-family, namely CXCL13 and CXCL14, are part of highly expressed ligand-receptor pairs (CXCL13-CXCR5 and CXCL14-CXCR4) in almost all immune aggregates except TLS number “5”. It might be an indication of exclusive biological processes undergoing in this TLS. All this information allowed us to classify some of the TLSs as myeloid enriched and mregDCs-enriched/depleted (**Figure 3E**).

### Gradient analysis of TLSs might be beneficial for understanding underlying biological processes

After describing 11 TLSs on one CRC tumor tissue, we realized that we were focused mainly on biological processes happening inside TLSs. To expand our understanding on TLSs’ microenvironment, we developed the “gradient analysis” algorithm which works in two steps. The first step is isolation of Visium spots which are equally distant from the border between the selected structure and neighboring spots, and either traversing towards the center (internal layers) or moving away from it (outside layers). The second step is regression of gene expression or cellular counts across the structure borderline, in our case from the TLS “core” towards the stroma/tumor region.

Applying this function to TLSs we were able to isolate 2 internal layers and 10 outside layers describing TLS microenvironment in the C2 CRC sample (**Figure 4A**). Then we were able to track gene expression changes across these layers. We call genes with gradual expression increase toward TLS “core” as “down” genes; genes with gradual increase outside TLSs --“up” genes. Thus, we extracted two gene signatures, “up” and “down”. “down” signature is composed of T cells (*TRBC1, TRBC2*), B cells (*CD79A, CD37*) and general inflammatory processes (*IKZF1*) markers. “up” signature mostly consists of stroma-related genes (*ACTG2, TGFB1L1*, **Figure 4B, Supplementary Figure S4**).

**Figure 4.**
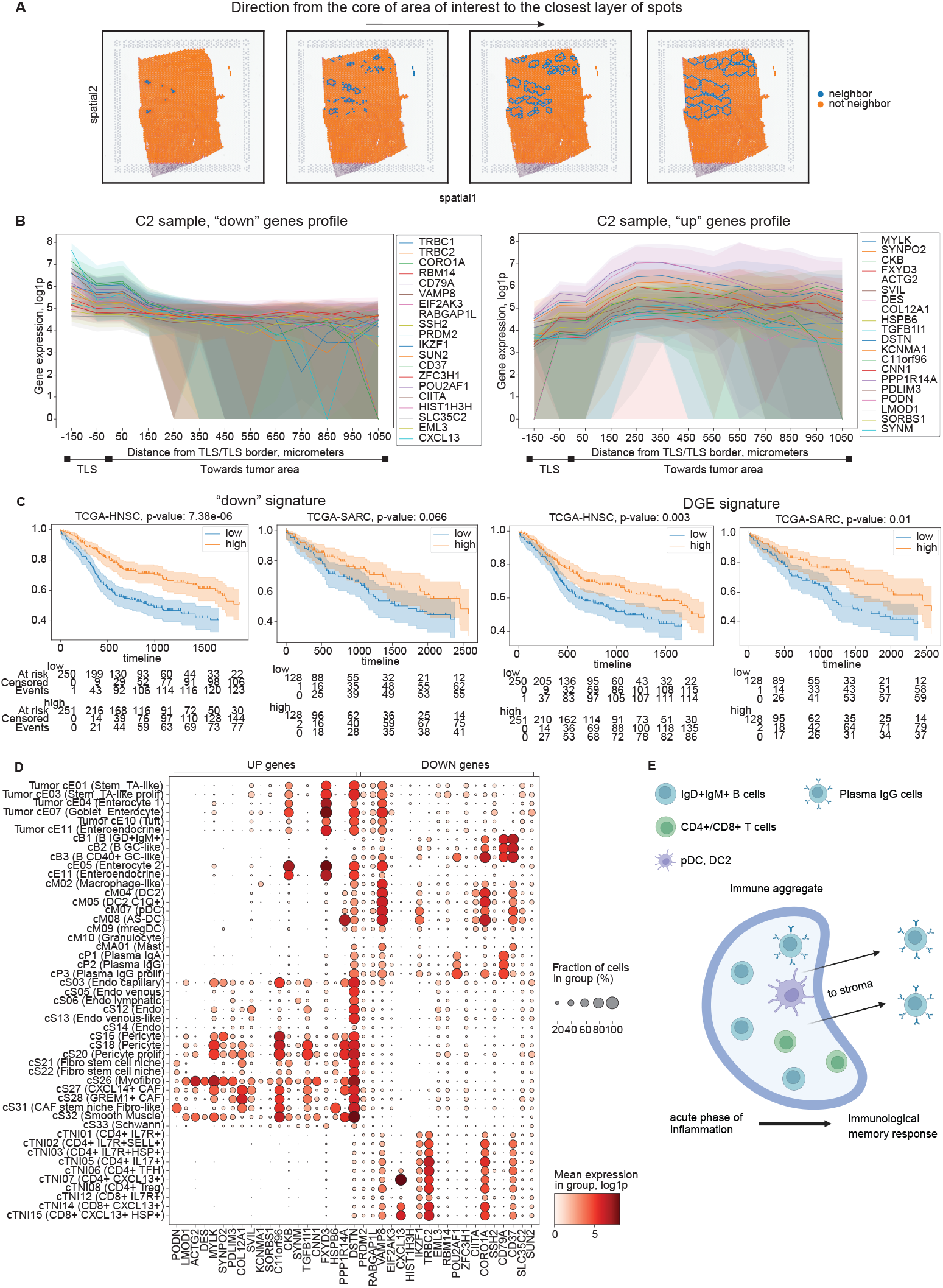
Gradient analysis of gene expression unveils gene signature different from differential gene expression signature. (A) Neighborhood analysis calculates spots coming from the core of the area of interest (immune aggregates or TLS in this case) toward the outside. Each step of the analysis adds a layer of spots at 100 micrometers from the previous layer. (B) Gene expression of regressed genes across neighborhoods at equally separated distances. Genes with positive or negative gradient (p-value <0.05) are selected. log1p is the natural logarithm of one plus the input array. (C) Kaplan-Meier curves reveal the difference between DOWN signature mined from gradient analysis across neighborhood layers (n=20) and signature mined from differential expression analysis (T-test, immune aggregates vs other areas, n=20) in the context of TCGA cancer datasets. (C) Only barcodes in neighborhood layers [n=1894] and genes that have non zero counts in more than 70 % of barcodes [n=1670] were used to find the signatures. DOWN signature genes: CD37, CD79A, CIITA, CORO1A, CXCL13, EIF2AK3, EML3, HIST1H3H, IKZF1, POU2AF1, PRDM2, RABGAP1L, RBM14, SLC35C2, SSH2, SUN2, TRBC1, TRBC2, VAMP8, ZFC3H1; differential gene expression signature genes: AKNA, CD37, CD53, CD74, CD79A, CIITA, CORO1A, CXCL13, CXCR4, IKZF1, ISG20, LAPTM5, LBH, POU2AF1, PTPN1, PTPRC, SMAP2, TRBC1, TRBC2, UBA52. Underscored genes are mutual for both signatures. (D) Dotplot of regressed genes’ expression in a colorectal cancer single-cell reference identifies cell types – source of gradiently expressed genes. (E) Cartoon depiction of major findings on immune aggregates with gene gradient analysis. Plasma IgG cells escape immune aggregates to the stroma region that highlights immunological memory response.

To assess clinical relevance of the collected signature, we performed survival analysis projecting signature activity on TCGA treatment naive bulk RNA-sequencing datasets (https://www.cancer.gov/tcga) using single sample gene set enrichment analysis (ssGSEA) approach ^28^. We compared obtained overall survival (OS) scores to the defined signature by performing differential gene expression analysis (DGE) comparing TLS zone against everything else. We observed similar power to predict patient cohorts with better survival (**Figure 4C**).

Finally, we developed a “back forward” approach to assign signature genes to the most probable cell type of origin defined by deconvolution model within each Visium spot annotated as TLS. As expected, genes identified to be gradually enriched in TLS zone were uniquely enriched in immune-related subsets while signature genes gradually downregulated in TLS were mainly detected in stroma/tumor cell types (**Figure 4D, Supplementary Figures S5, S6**). Taken together, the approach revealed the potential trafficking of plasma IgG cells from the TLS “core” toward outside TLSs in agreement with previously established role in immunological response ^11,29^ (**Figure 4E**; figure was created with BioRender.com).

We also combined two described approaches – splitting immune aggregates into separate entities and gradient analysis – to assess differences in the closest microenvironment between TLSs in the C2 CRC sample (**Supplementary Figure S7**). In addition to what we have already shown on **Figure 3E**, where we classified some of the TLSs into myeloid enriched and mregDC-enriched/depleted categories, we also discovered additional differences in the closest microenvironment. For example, the neighborhood of TLS “0” is enriched in *IGHG4* and *IGLV4-60* (**Supplementary Figure S7**). We were not able to classify cluster “0” into one of the above categories before, but with the help of the combined approach it may give an insight into underlying biological mechanisms happening not only in TLS but in its neighborhood as well.

## DISCUSSION

In this study we developed a new methodology to analyze spatial transcriptomics data. We proposed a standalone Python-based pipeline which combines pathologist-driven slide annotation, projection of cell populations within each Visium spot with absolute quantification based on accurate nuclei segmentation. Also, we developed novel functions performing pattern identification within clustering results of cellular compositions, automated segmentation of geographically separated domains of similar pattern, statistical framework to compare compositions of different domains and toolkit perform gradient analysis: regression of cellular quantities over the borderline of region of interest.

Due to sample size limitations, we cannot properly benchmark our toolkit, however using colorectal C2 sample as a proof-of-concept we reassure it properly reflects the underlying biology and describes tumor microenvironment components and distributions. With unsupervised clustering of cellular compositions, we identified patterns matching perfectly supervised pathological annotation and aligned with expected biological pathway activities enriched in those areas (**Figure 1**). Later we zoomed into TLS areas and delineated the immunological reactions known to be spatially organized, completely in an unsupervised manner (**Figure 2**). Besides from the validation of certain biological patterns we believe our toolkit can serve exploratory goals as we were able to classify TLS subtypes using developed statistical framework (**Figure 3**). We also extracted gradient signature of gene expression that gradually increased (or decreased) relative to the TLS border line. These signatures proved to be predictive in treatment naive setting of TCGA cohorts. For example, analysis performed on head-and-neck cancer (TCGA-HNSC) shows statistically significant difference in overall survival between patients with “low” and “high” TLS signatures scores. Similar separation was observed in the context of sarcoma (TCGA-SARC) (**Figure 4**). These survival analysis results are in line with previously published articles stating that immune infiltration is survival-promoting in head-and-neck cancer ^30^ and sarcoma contexts ^31^.

In conclusion, spatial transcriptomics represents a paradigm shift in our approach to dissecting the complexities of gene expression within the spatial context of tissues. Thus, engineering new algorithms and corresponding software is essential to unlock biological discovery. Its integration into biological research promises to unravel fundamental biological principles, accelerate biomedical discoveries, and ultimately transform our understanding and treatment of human health and disease.

## MATERIALS AND METHODS

### Tested samples

The tested samples included endometrium (n=21), melanoma (n=2), lung (n=5) and colorectal (n=7) human cancers for benchmarking purposes. Three colorectal samples were used for the subsequent analysis. All the tested samples were stained using hematoxylin and eosin. Visium transcriptome data was obtained using standard procedure provided by 10X Genomics for FFPE tissue sections (https://www.10xgenomics.com/products/spatial-gene-expression).

### Data sharing statement

All raw and processed data is available upon request at CIDC-CIMAC portal: https://cidc.nci.nih.gov.

### Deconvolution benchmark

For the deconvolution step we tested a set of publicly available algorithms: Cell2location, Stereoscope, RCTD, Seurat, Tangram, SpatialDWLS, DestVI.used for deconvolution included single cell datasets available in-house. For the benchmark we contrasted results of deconvolutions to pathology supervised annotations performed in QuPath ^32^. Alternatively, we used an image-based classifier implemented in QuPath to classify segmented cells into tumor, immune or stroma types. The quantities of these three classes per aligned Visium spot were used as ground truth to compare deconvolution results as well. Statistical analysis was performed using scikit implementation of ROC analysis ^33^.

### Pathway analysis

Pathway analysis was performed using decoupleR implementation of multivariate linear model, over-representation or gene set enrichment analysis ^34^. Databases listed in **Figure 1 –** PROGENy, DoRothEA, Reactome – were used to calculate signature enrichments in Visium samples. We used GSEApy ^35^ implementation of gene set enrichment analysis for analysis of separate immune aggregates. Results are provided in **Supplementary Figures S4, S7**. Used databases – GO Cellular Component ^36^, KEGG ^37^, LINCS ^38^, MSigDB ^39^ and Reactome.

### Receptor-ligand interaction analysis

The analysis was performed using squidpy implementation of permutation test as described in CellPhoneDB original paper ^40^.

### Gradient analysis

Gradient analysis is performed in two steps. The first step is calculating neighboring layers of Visium spots to a selected tissue region (**Figure 4A**). The algorithm distinguishes between internal layers (spots inside the selected structure) and outer layers (spots outside the selected structure). Since neighboring layers are selected, the algorithm can regress gene expression or cellular counts traversing across the structure borderline (**Figure 4B**). It is the second step. By default, it is done using statsmodels implementation of generalized linear models ^41^.

### Survival analysis

Survival analysis was performed on TCGA bulk RNA-sequencing cohorts (n=33) [https://www.cancer.gov/tcga]. The Python implementation of Kaplan-Meier estimate function was used from lifelines package ^42^.

### Code sharing statement

All the code used for cell segmentation and Visium spot deconvolution is available at Human Immune Monitoring Center (HIMC) GitHub page https://github.com/ismms-himc/visium_segmentation, https://github.com/ismms-himc/visium_deconvolution. The downstream analysis part is also accessible via HIMC GitHub page as an installable Python package https://github.com/ismms-himc/Visium_analysis. Additional questions regarding analysis can be addressed via email to.

### Software and package versions

All the necessary Python packages used in the cell segmentation, Visium spot deconvolution and the downstream analysis are listed in python or configuration files at the projects’ GitHub pages (see “Code sharing statement” for the GitHub links).

## ACKNOWLEDGEMENT

We are grateful to Boehringer Ingelheim Pharmaceuticals for their sponsorship and support in providing resources for sample collection.

This work was supported in part through the computational and data resources and staff expertise provided by Scientific Computing and Data at the Icahn School of Medicine at Mount Sinai and supported by the Clinical and Translational Science Awards (CTSA) grant UL1TR004419 from the National Center for Advancing Translational Sciences.

## Notes

### Competing Interest Statement

The authors have declared no competing interest.

