## Supplementary figures for "Informing biologically relevant signal from spatial transcriptomic data"

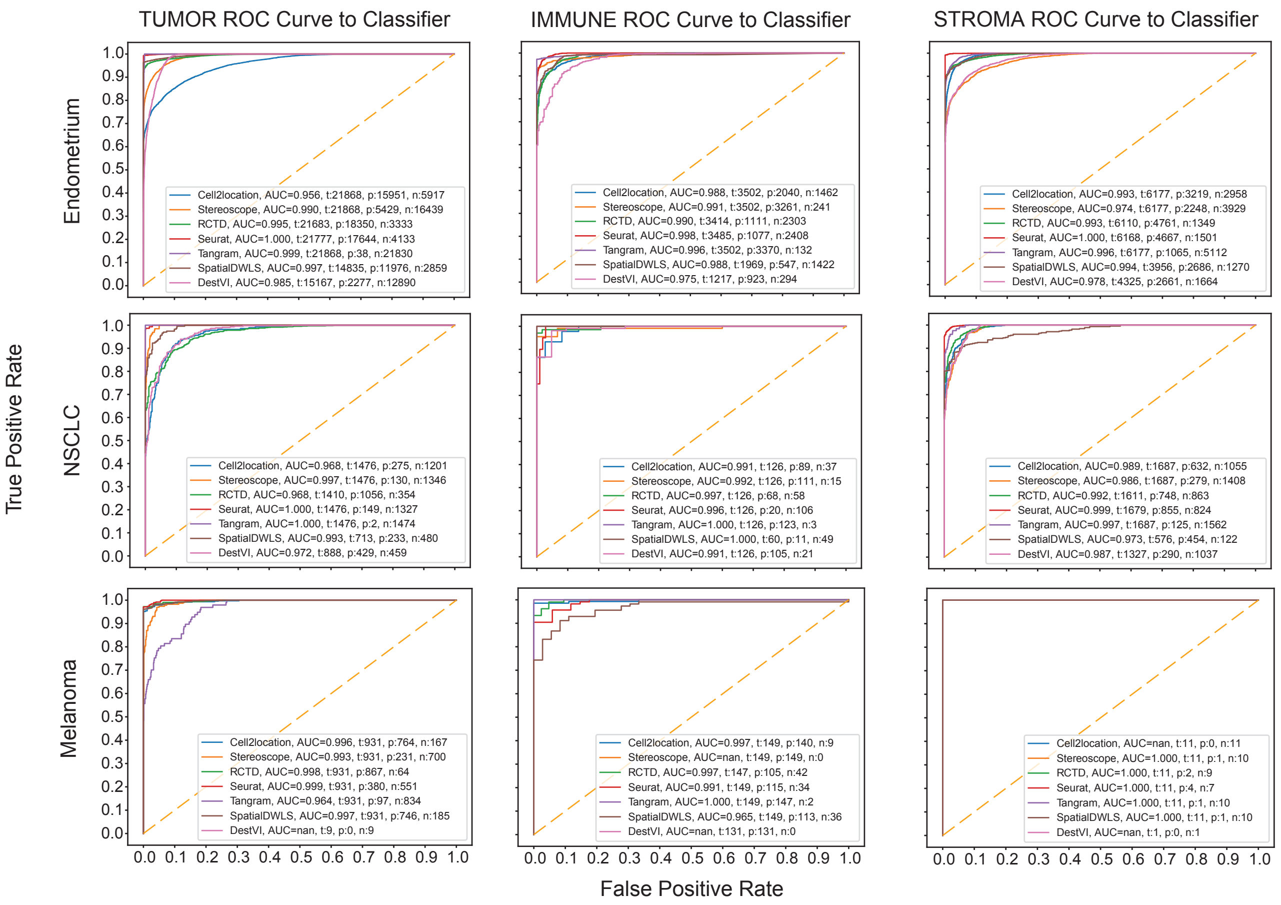

TUMOR ROC Curve to Classifier
IMMUNE ROC Curve to Classifier
STROMA ROC Curve to Classifier
Endometrium
NSCLC
Melanoma
True Positive Rate
False Positive Rate
Cell2location, AUC=0.956, t:21868, p:15951, n:5917
Stereoscope, AUC=0.990, t:21868, p:5429, n:16439
RCTD, AUC=0.995, t:21683, p:18350, n:3333
Seurat, AUC=1.000, t:21777, p:17644, n:4133
Tangram, AUC=0.999, t:21868, p:38, n:21830
SpatialDWLS, AUC=0.997, t:14835, p:11976, n:2859
DestVI, AUC=0.985, t:15167, p:2277, n:12890
Cell2location, AUC=0.988, t:3502, p:2040, n:1462
Stereoscope, AUC=0.991, t:3502, p:3261, n:241
RCTD, AUC=0.990, t:3414, p:1111, n:2303
Seurat, AUC=0.998, t:3485, p:1077, n:2408
Tangram, AUC=0.996, t:3502, p:3370, n:132
SpatialDWLS, AUC=0.988, t:1969, p:547, n:1422
DestVI, AUC=0.975, t:1217, p:923, n:294
Cell2location, AUC=0.993, t:6177, p:3219, n:2958
Stereoscope, AUC=0.974, t:6177, p:2248, n:3929
RCTD, AUC=0.993, t:6110, p:4761, n:1349
Seurat, AUC=1.000, t:6168, p:4667, n:1501
Tangram, AUC=0.996, t:6177, p:1065, n:5112
SpatialDWLS, AUC=0.994, t:3956, p:2686, n:1270
DestVI, AUC=0.978, t:4325, p:2661, n:1664
Cell2location, AUC=0.968, t:1476, p:275, n:1201
Stereoscope, AUC=0.997, t:1476, p:130, n:1346
RCTD, AUC=0.968, t:1410, p:1056, n:354
Seurat, AUC=1.000, t:1476, p:149, n:1327
Tangram, AUC=1.000, t:1476, p:2, n:1474
SpatialDWLS, AUC=0.993, t:713, p:233, n:480
DestVI, AUC=0.972, t:888, p:429, n:459
Cell2location, AUC=0.991, t:126, p:89, n:37
Stereoscope, AUC=0.992, t:126, p:111, n:15
RCTD, AUC=0.997, t:126, p:68, n:58
Seurat, AUC=0.996, t:126, p:20, n:106
Tangram, AUC=1.000, t:126, p:123, n:3
SpatialDWLS, AUC=1.000, t:60, p:11, n:49
DestVI, AUC=0.991, t:126, p:105, n:21
Cell2location, AUC=0.989, t:1687, p:632, n:1055
Stereoscope, AUC=0.986, t:1687, p:279, n:1408
RCTD, AUC=0.992, t:1611, p:748, n:863
Seurat, AUC=0.999, t:1679, p:855, n:824
Tangram, AUC=0.997, t:1687, p:125, n:1562
SpatialDWLS, AUC=0.973, t:576, p:454, n:122
DestVI, AUC=0.987, t:1327, p:290, n:1037
Cell2location, AUC=0.996, t:931, p:764, n:167
Stereoscope, AUC=0.993, t:931, p:231, n:700
RCTD, AUC=0.998, t:931, p:867, n:64
Seurat, AUC=0.999, t:931, p:380, n:551
Tangram, AUC=0.964, t:931, p:97, n:834
SpatialDWLS, AUC=0.997, t:931, p:746, n:185
DestVI, AUC=nan, t:9, p:0, n:9
Cell2location, AUC=0.997, t:149, p:140, n:9
Stereoscope, AUC=nan, t:149, p:149, n:0
RCTD, AUC=0.997, t:147, p:105, n:42
Seurat, AUC=0.991, t:149, p:115, n:34
Tangram, AUC=1.000, t:149, p:147, n:2
SpatialDWLS, AUC=0.965, t:149, p:113, n:36
DestVI, AUC=nan, t:131, p:131, n:0
Cell2location, AUC=nan, t:11, p:0, n:11
Stereoscope, AUC=1.000, t:11, p:1, n:10
RCTD, AUC=1.000, t:11, p:2, n:9
Seurat, AUC=1.000, t:11, p:4, n:7
Tangram, AUC=1.000, t:11, p:1, n:10
SpatialDWLS, AUC=1.000, t:11, p:1, n:10
DestVI, AUC=nan, t:1, p:0, n:1

**Figure S1 (previous page). Benchmarking of various RNA spot-based deconvolution tools in a variety of diseased tissues. ROC curves and AUC scores.**
Receiver Operating Characteristic Curve (ROC) curves are plotted for each tool within a pathological area of a tissue from which Area Under the Curve (AUC) scores were subsequently calculated. Deconvolution outputs were compared against ground truth spot labels expertly annotated by a trained pathologist. Ground truth labels were classified as either Tumor, Immune, Stroma or Other. Legend of the ROC curves show for each tool the AUC score, total spots (t), positive spots (p), meaning labels that were concordant between the deconvolution tool and pathologist and vice versa with negative spots (n).

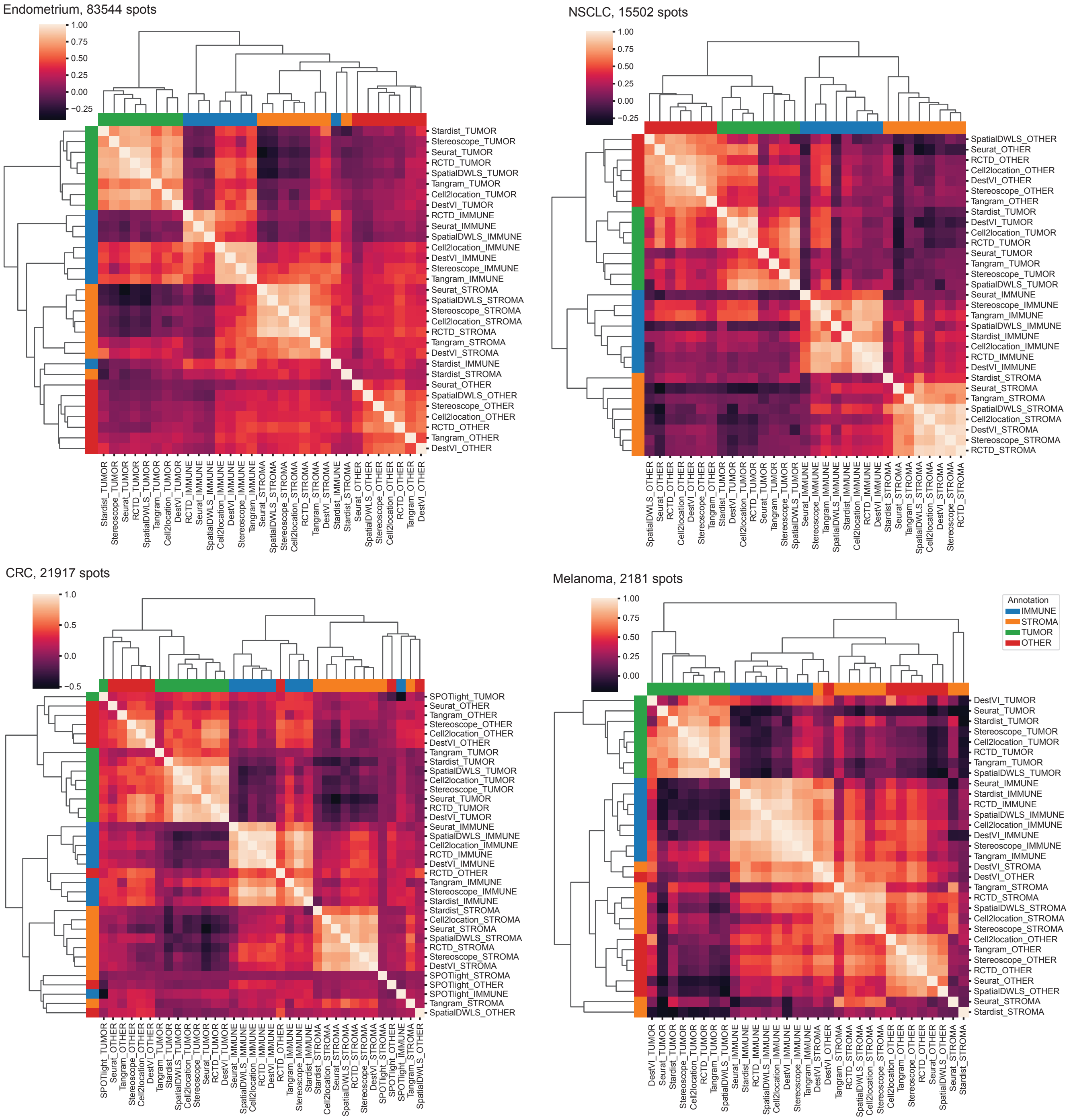
Endometrium, 83544 spots
1.00
0.75
0.50
0.25
0.00
−0.25
Stardist_TUMOR
Stereoscope_TUMOR
Seurat_TUMOR
RCTD_TUMOR
SpatialDWLS_TUMOR
Tangram_TUMOR
Cell2location_TUMOR
DestVI_TUMOR
RCTD_IMMUNE
Seurat_IMMUNE
SpatialDWLS_IMMUNE
Cell2location_IMMUNE
DestVI_IMMUNE
Stereoscope_IMMUNE
Tangram_IMMUNE
Seurat_STROMA
SpatialDWLS_STROMA
Stereoscope_STROMA
Cell2location_STROMA
RCTD_STROMA
Tangram_STROMA
DestVI_STROMA
Stardist_IMMUNE
Stardist_STROMA
Seurat_OTHER
SpatialDWLS_OTHER
Stereoscope_OTHER
Cell2location_OTHER
RCTD_OTHER
Tangram_OTHER
DestVI_OTHER
NSCLC, 15502 spots
1.00
0.75
0.50
0.25
0.00
−0.25
SpatialDWLS_OTHER
Seurat_OTHER
RCTD_OTHER
Cell2location_OTHER
DestVI_OTHER
Stereoscope_OTHER
Tangram_OTHER
Stardist_TUMOR
DestVI_TUMOR
Cell2location_TUMOR
RCTD_TUMOR
Seurat_TUMOR
Tangram_TUMOR
Stereoscope_TUMOR
SpatialDWLS_TUMOR
Seurat_IMMUNE
Stereoscope_IMMUNE
Tangram_IMMUNE
SpatialDWLS_IMMUNE
Stardist_IMMUNE
Cell2location_IMMUNE
RCTD_IMMUNE
DestVI_IMMUNE
Stardist_STROMA
Seurat_STROMA
Tangram_STROMA
SpatialDWLS_STROMA
Cell2location_STROMA
DestVI_STROMA
Stereoscope_STROMA
RCTD_STROMA
CRC, 21917 spots
1.0
0.5
0.0
−0.5
SPOTlight_TUMOR
Seurat_OTHER
Tangram_OTHER
Stereoscope_OTHER
Cell2location_OTHER
DestVI_OTHER
Tangram_TUMOR
Stardist_TUMOR
SpatialDWLS_TUMOR
Cell2location_TUMOR
Stereoscope_TUMOR
Seurat_TUMOR
RCTD_TUMOR
DestVI_TUMOR
Seurat_IMMUNE
SpatialDWLS_IMMUNE
Cell2location_IMMUNE
RCTD_IMMUNE
DestVI_IMMUNE
RCTD_OTHER
Tangram_IMMUNE
Stereoscope_IMMUNE
Stardist_IMMUNE
Stardist_STROMA
Cell2location_STROMA
Seurat_STROMA
SpatialDWLS_STROMA
RCTD_STROMA
Stereoscope_STROMA
DestVI_STROMA
SPOTlight_STROMA
SPOTlight_OTHER
SPOTlight_IMMUNE
Tangram_STROMA
SpatialDWLS_OTHER
Melanoma, 2181 spots
1.00
0.75
0.50
0.25
0.00
Annotation
IMMUNE
STROMA
TUMOR
OTHER
DestVI_TUMOR
Seurat_TUMOR
Stardist_TUMOR
Stereoscope_TUMOR
Cell2location_TUMOR
RCTD_TUMOR
Tangram_TUMOR
SpatialDWLS_TUMOR
Seurat_IMMUNE
Stardist_IMMUNE
RCTD_IMMUNE
SpatialDWLS_IMMUNE
Cell2location_IMMUNE
DestVI_IMMUNE
Stereoscope_IMMUNE
Tangram_IMMUNE
DestVI_STROMA
DestVI_OTHER
Tangram_STROMA
RCTD_STROMA
SpatialDWLS_STROMA
Cell2location_STROMA
Stereoscope_STROMA
Cell2location_OTHER
Tangram_OTHER
Stereoscope_OTHER
RCTD_OTHER
Seurat_OTHER
SpatialDWLS_OTHER
Seurat_STROMA
Stardist_STROMA

**Figure S2 (previous page). Benchmarking of various RNA spot-based deconvolution tools in a variety of diseased tissues. Correlation heatmaps.**

Heatmaps show the correlation matrix of total cells per spot between each tool in a pathological area. All tools show good agreement between each other no matter the cell or tissue type.

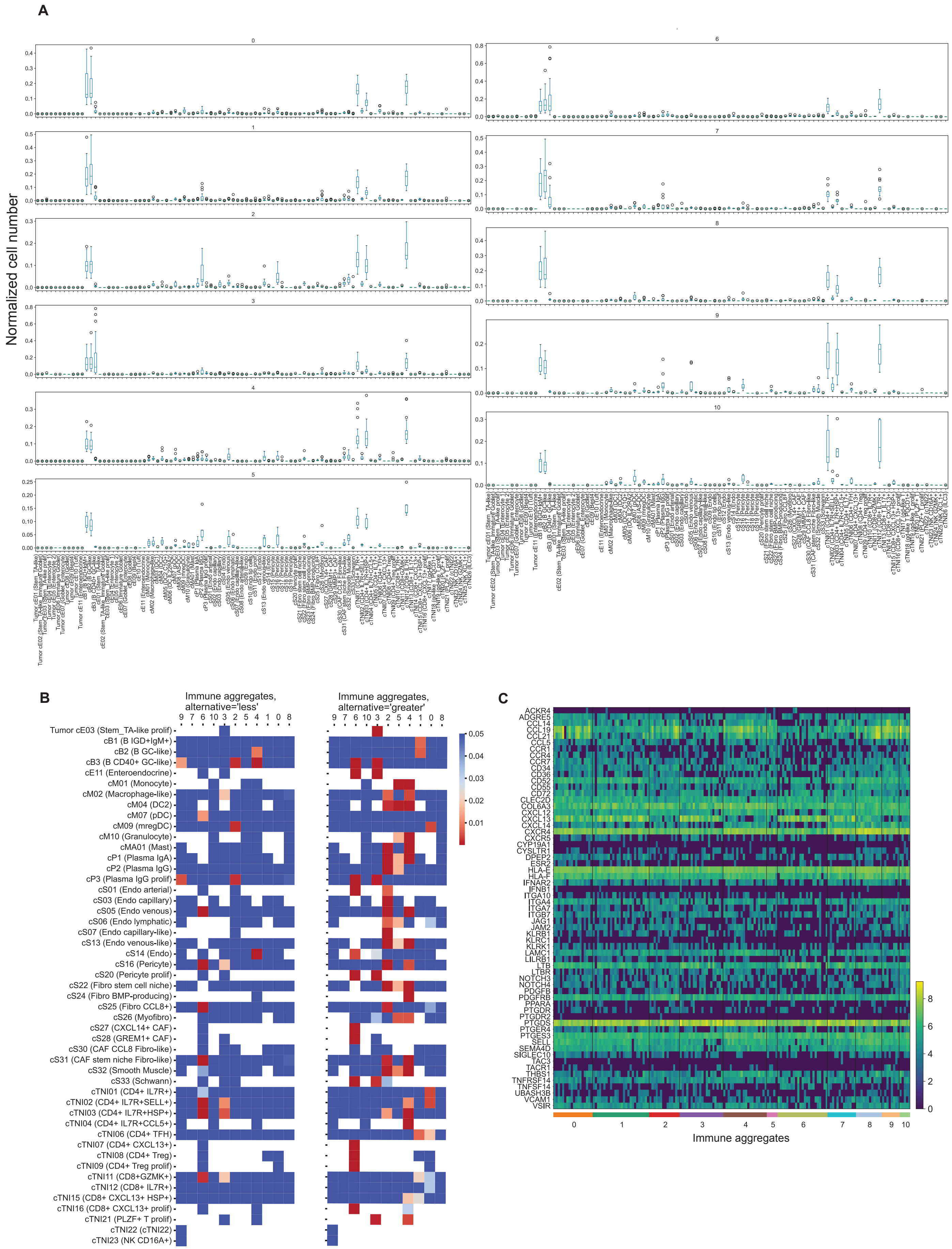
A
Normalized cell number
B
Immune aggregates, alternative='less'
Immune aggregates, alternative='greater'
9 7 6 10 3 2 5 4 1 0 8
Tumor cE03 (Stem_TA-like prolif)
cB1 (B IGD+IgM+)
cB2 (B GC-like)
cB3 (B CD40+ GC-like)
cE11 (Enteroendocrine)
cM01 (Monocyte)
cM02 (Macrophage-like)
cM04 (DC2)
cM07 (pDC)
cM09 (mregDC)
cM10 (Granulocyte)
cMA01 (Mast)
cP1 (Plasma IgA)
cP2 (Plasma IgG)
cP3 (Plasma IgG prolif)
cS01 (Endo arterial)
cS03 (Endo capillary)
cS05 (Endo venous)
cS06 (Endo lymphatic)
cS07 (Endo capillary-like)
cS13 (Endo venous-like)
cS14 (Endo)
cS16 (Pericyte)
cS20 (Pericyte prolif)
cS22 (Fibro stem cell niche)
cS24 (Fibro BMP-producing)
cS25 (Fibro CCL8+)
cS26 (Myofibro)
cS27 (CXCL14+ CAF)
cS28 (GREM1+ CAF)
cS30 (CAF CCL8 Fibro-like)
cS31 (CAF stem niche Fibro-like)
cS32 (Smooth Muscle)
cS33 (Schwann)
cTNI01 (CD4+ IL7R+)
cTNI02 (CD4+ IL7R+SELL+)
cTNI03 (CD4+ IL7R+HSP+)
cTNI04 (CD4+ IL7R+CCL5+)
cTNI06 (CD4+ TFH)
cTNI07 (CD4+ CXCL13+)
cTNI08 (CD4+ Treg)
cTNI09 (CD4+ Treg prolif)
cTNI11 (CD8+GZMK+)
cTNI12 (CD8+ IL7R+)
cTNI15 (CD8+ CXCL13+ HSP+)
cTNI16 (CD8+ CXCL13+ prolif)
cTNI21 (PLZF+ T prolif)
cTNI22 (cTNI22)
cTNI23 (NK CD16A+)
0.05
0.04
0.03
0.02
0.01
C
ACKR4
ADGRE5
CCL14
CCL19
CCL21
CCL5
CCR1
CCR4
CCR7
CD34
CD36
CD52
CD55
CD72
CLEC2D
COL6A3
CXCL12
CXCL13
CXCL14
CXCR4
CXCR5
CYP19A1
CYSLTR1
DPEP2
ESR2
HLA-E
HLA-F
IFNAR2
IFNB1
ITGA10
ITGA4
ITGA7
ITGB7
JAG1
JAM2
KLRB1
KLRC1
KLRK1
LAMC1
LILRB1
LTB
LTBR
NOTCH3
NOTCH4
PDGFB
PDGFRB
PPARA
PTGDR
PTGDR2
PTGDS
PTGER4
PTGES3
SELL
SEMA4D
SIGLEC10
TAC3
TACR1
THBS1
TNFRSF14
TNFSF14
UBASH3B
VCAM1
VSIR
8
6
4
2
0
0 1 2 3 4 5 6 7 8 9 10
Immune aggregates

**Figure S3 (previous page). Characteristics of separate immune aggregates on C2 (see Figure 2A) histological slide.**

(A) Distribution of normalized cell number per aggregate across all cell types. (B) Heatmap of corrected p-values testing one immune aggregate vs other aggregates (Mann-Whitney U test, Benjamini-Hochberg multiple correction). (C) Heatmap of gene expression of ligands and receptors listed in (see Figure 3D) across the immune aggregates.

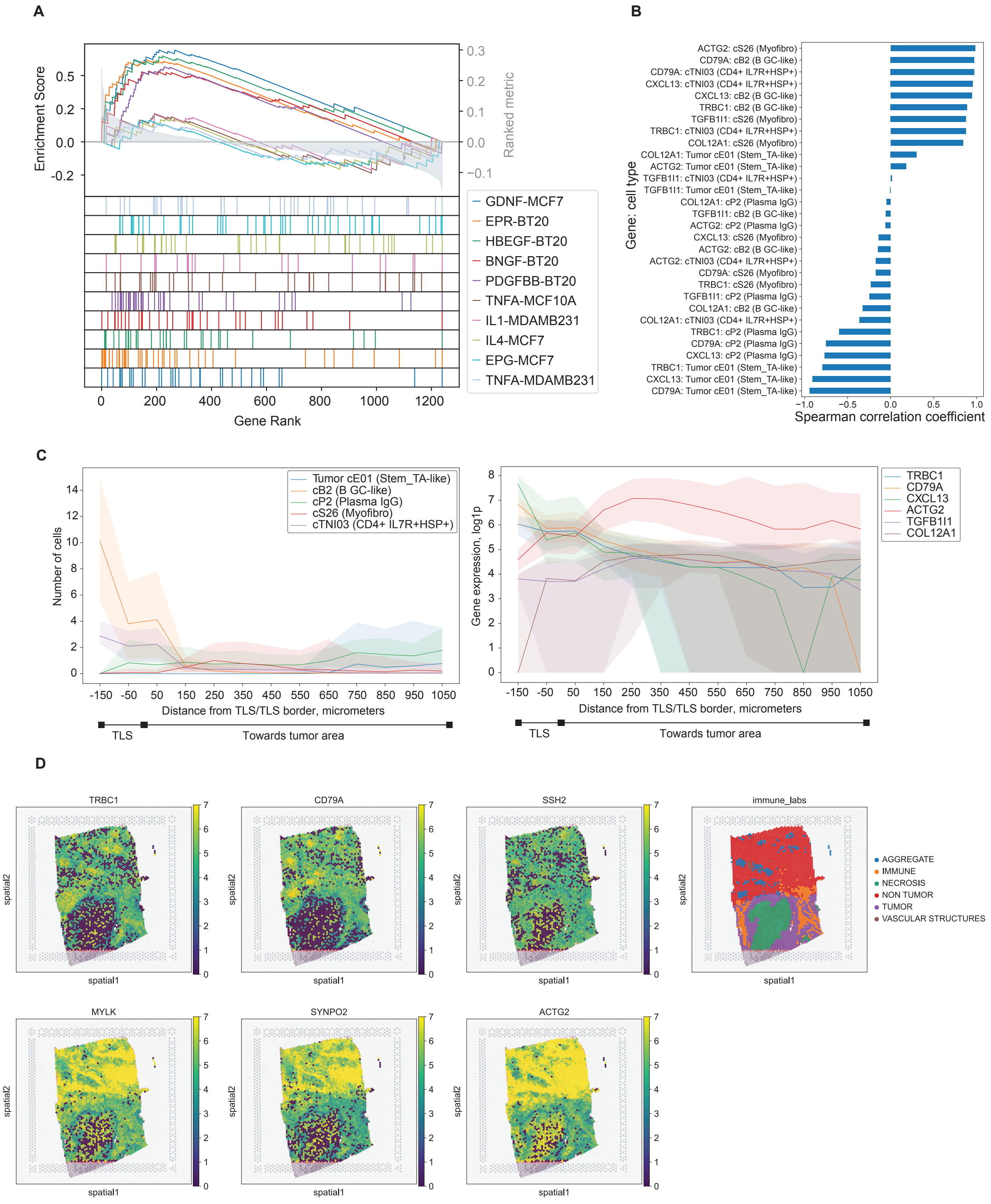
A
Enrichment Score
Ranked metric
Gene Rank
GDNF-MCF7
EPR-BT20
HBEGF-BT20
BNGF-BT20
PDGFBB-BT20
TNFA-MCF10A
IL1-MDAMB231
IL4-MCF7
EPG-MCF7
TNFA-MDAMB231
B
Gene: cell type
ACTG2: cS26 (Myofibro)
CD79A: cB2 (B GC-like)
CD79A: cTNI03 (CD4+ IL7R+HSP+)
CXCL13: cTNI03 (CD4+ IL7R+HSP+)
CXCL13: cB2 (B GC-like)
TRBC1: cB2 (B GC-like)
TGFB1I1: cS26 (Myofibro)
TRBC1: cTNI03 (CD4+ IL7R+HSP+)
COL12A1: cS26 (Myofibro)
COL12A1: Tumor cE01 (Stem_TA-like)
ACTG2: Tumor cE01 (Stem_TA-like)
TGFB1I1: cTNI03 (CD4+ IL7R+HSP+)
TGFB1I1: Tumor cE01 (Stem_TA-like)
COL12A1: cP2 (Plasma IgG)
TGFB1I1: cB2 (B GC-like)
ACTG2: cP2 (Plasma IgG)
CXCL13: cS26 (Myofibro)
ACTG2: cB2 (B GC-like)
ACTG2: cTNI03 (CD4+ IL7R+HSP+)
CD79A: cS26 (Myofibro)
TRBC1: cS26 (Myofibro)
TGFB1I1: cP2 (Plasma IgG)
COL12A1: cB2 (B GC-like)
COL12A1: cTNI03 (CD4+ IL7R+HSP+)
TRBC1: cP2 (Plasma IgG)
CD79A: cP2 (Plasma IgG)
CXCL13: cP2 (Plasma IgG)
TRBC1: Tumor cE01 (Stem_TA-like)
CXCL13: Tumor cE01 (Stem_TA-like)
CD79A: Tumor cE01 (Stem_TA-like)
Spearman correlation coefficient
C
Number of cells
Tumor cE01 (Stem_TA-like)
cB2 (B GC-like)
cP2 (Plasma IgG)
cS26 (Myofibro)
cTNI03 (CD4+ IL7R+HSP+)
Distance from TLS/TLS border, micrometers
TLS
Towards tumor area
Gene expression, log1p
TRBC1
CD79A
CXCL13
ACTG2
TGFB1I1
COL12A1
Distance from TLS/TLS border, micrometers
TLS
Towards tumor area
D
TRBC1
CD79A
SSH2
immune_labs
AGGREGATE
IMMUNE
NECROSIS
NON TUMOR
TUMOR
VASCULAR STRUCTURES
MYLK
SYNPO2
ACTG2
spatial1
spatial2

**Figure S4 (previous page). Additional characteristics of regressed genes.**
(A) Gene set enrichment analysis results, where top left curves represent gene signatures enriched in the immune aggregate area and bottom right curves represent gene signatures enriched in the neighborhood of immune aggregate area. All depicted signatures were taken from the LINCS database. (B) Spearman correlation coefficients between "down"/"up" genes and cell types where they are most abundant. (C) Distribution of genes and cell types from Supplementary FigureS3B across the layers. (D) Example of expression of "down" genes (top panel) and "up" genes (bottom panel).

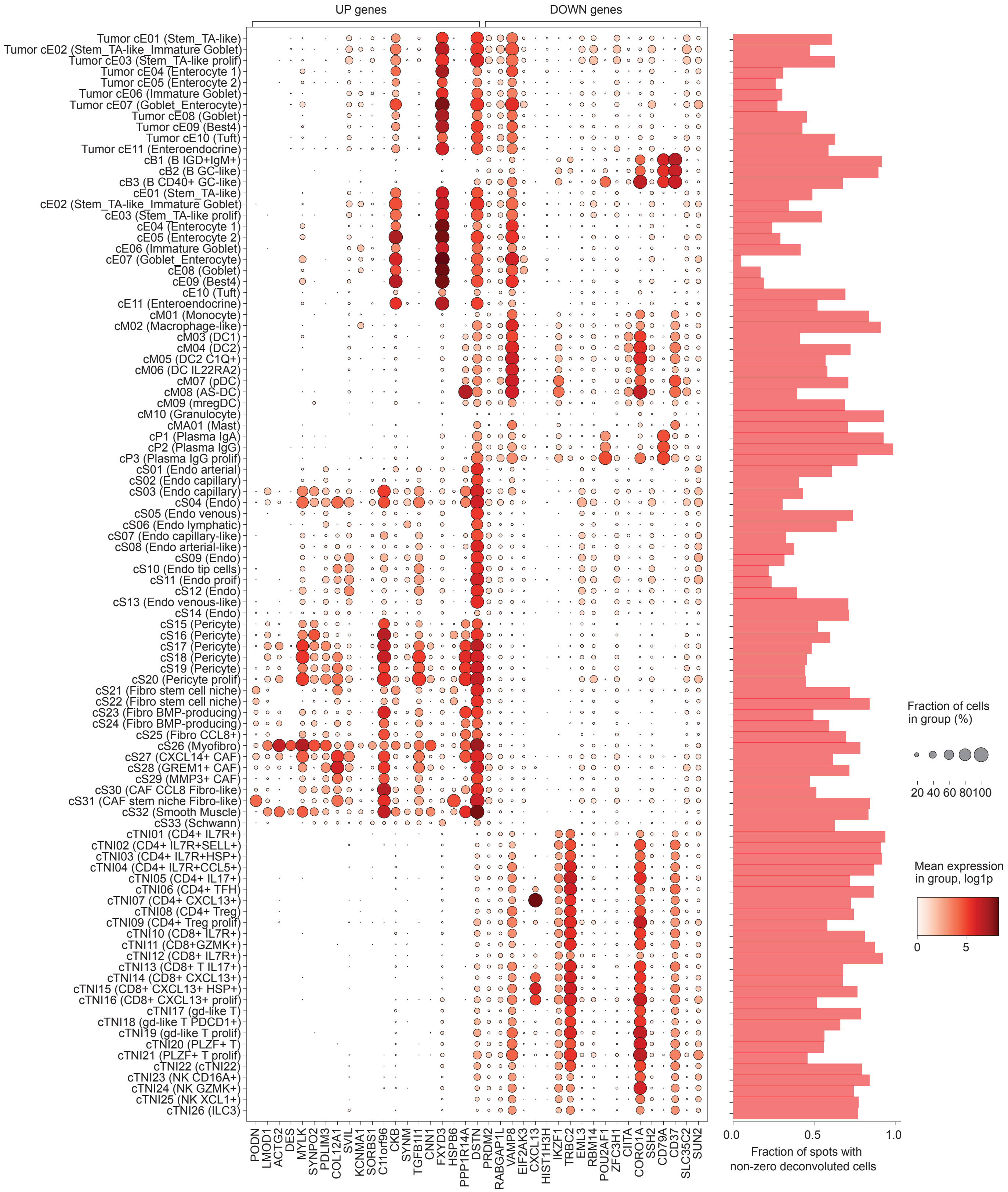

UP genes
DOWN genes
Tumor cE01 (Stem_TA-like)
Tumor cE02 (Stem_TA-like_Immature Goblet)
Tumor cE03 (Stem_TA-like prolif)
Tumor cE04 (Enterocyte 1)
Tumor cE05 (Enterocyte 2)
Tumor cE06 (Immature Goblet)
Tumor cE07 (Goblet_Enterocyte)
Tumor cE08 (Goblet)
Tumor cE09 (Best4)
Tumor cE10 (Tuft)
Tumor cE11 (Enteroendocrine)
cB1 (B IGD+IgM+)
cB2 (B GC-like)
cB3 (B CD40+ GC-like)
cE01 (Stem_TA-like)
cE02 (Stem_TA-like_Immature Goblet)
cE03 (Stem_TA-like prolif)
cE04 (Enterocyte 1)
cE05 (Enterocyte 2)
cE06 (Immature Goblet)
cE07 (Goblet_Enterocyte)
cE08 (Goblet)
cE09 (Best4)
cE10 (Tuft)
cE11 (Enteroendocrine)
cM01 (Monocyte)
cM02 (Macrophage-like)
cM03 (DC1)
cM04 (DC2)
cM05 (DC2 C1Q+)
cM06 (DC IL22RA2)
cM07 (pDC)
cM08 (AS-DC)
cM09 (mregDC)
cM10 (Granulocyte)
cMA01 (Mast)
cP1 (Plasma IgA)
cP2 (Plasma IgG)
cP3 (Plasma IgG prolif)
cS01 (Endo arterial)
cS02 (Endo capillary)
cS03 (Endo capillary)
cS04 (Endo)
cS05 (Endo venous)
cS06 (Endo lymphatic)
cS07 (Endo capillary-like)
cS08 (Endo arterial-like)
cS09 (Endo)
cS10 (Endo tip cells)
cS11 (Endo proif)
cS12 (Endo)
cS13 (Endo venous-like)
cS14 (Endo)
cS15 (Pericyte)
cS16 (Pericyte)
cS17 (Pericyte)
cS18 (Pericyte)
cS19 (Pericyte)
cS20 (Pericyte prolif)
cS21 (Fibro stem cell niche)
cS22 (Fibro stem cell niche)
cS23 (Fibro BMP-producing)
cS24 (Fibro BMP-producing)
cS25 (Fibro CCL8+)
cS26 (Myofibro)
cS27 (CXCL14+ CAF)
cS28 (GREM1+ CAF)
cS29 (MMP3+ CAF)
cS30 (CAF CCL8 Fibro-like)
cS31 (CAF stem niche Fibro-like)
cS32 (Smooth Muscle)
cS33 (Schwann)
cTNI01 (CD4+ IL7R+)
cTNI02 (CD4+ IL7R+SELL+)
cTNI03 (CD4+ IL7R+HSP+)
cTNI04 (CD4+ IL7R+CCL5+)
cTNI05 (CD4+ IL17+)
cTNI06 (CD4+ TFH)
cTNI07 (CD4+ CXCL13+)
cTNI08 (CD4+ Treg)
cTNI09 (CD4+ Treg prolif)
cTNI10 (CD8+ IL7R+)
cTNI11 (CD8+GZMK+)
cTNI12 (CD8+ IL7R+)
cTNI13 (CD8+ T IL17+)
cTNI14 (CD8+ CXCL13+)
cTNI15 (CD8+ CXCL13+ HSP+)
cTNI16 (CD8+ CXCL13+ prolif)
cTNI17 (gd-like T)
cTNI18 (gd-like T PDCD1+)
cTNI19 (gd-like T prolif)
cTNI20 (PLZF+ T)
cTNI21 (PLZF+ T prolif)
cTNI22 (cTNI22)
cTNI23 (NK CD16A+)
cTNI24 (NK GZMK+)
cTNI25 (NK XCL1+)
cTNI26 (ILC3)
PODN
LMOD1
ACTG2
DES
MYLK
SYNPO2
PDLIM3
COL12A1
SVIL
KCNMA1
SORBS1
C11orf96
CKB
SYNM
TGFB1I1
CNN1
FXYD3
HSPB6
PPP1R14A
DSTN
PRDM2
RABGAP1L
VAMP8
EIF2AK3
CXCL13
HIST1H3H
IKZF1
TRBC2
EML3
RBM14
POU2AF1
ZFC3H1
CIITA
CORO1A
SSH2
CD79A
CD37
SLC35C2
SUN2
0.0
0.5
1.0
Fraction of spots with
non-zero deconvoluted cells
Fraction of cells
in group (%)
20 40 60 80100
Mean expression
in group, log1p
0
5

**Figure S5 (previous page). Dotplot of regressed genes' expression in a colorectal cancer single-cell reference (Pelka et al, 2021) with a right panel showing fraction of spots with non-zero deconvoluted cells on a histological slide.**

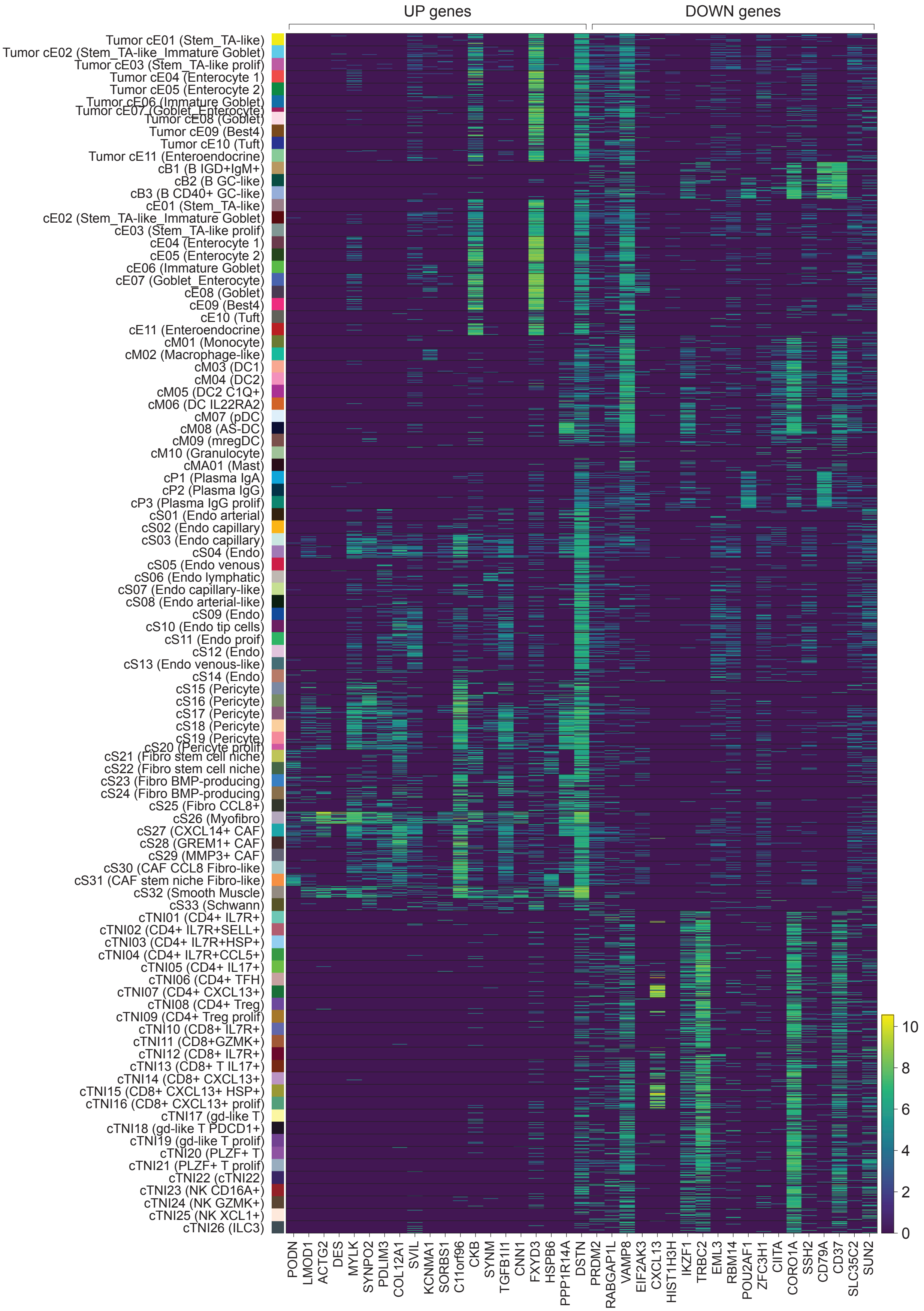

UP genes
DOWN genes
Tumor cE01 (Stem_TA-like)
Tumor cE02 (Stem_TA-like_Immature Goblet)
Tumor cE03 (Stem_TA-like prolif)
Tumor cE04 (Enterocyte 1)
Tumor cE05 (Enterocyte 2)
Tumor cE06 (Immature Goblet)
Tumor cE07 (Goblet_Enterocyte)
Tumor cE08 (Goblet)
Tumor cE09 (Best4)
Tumor cE10 (Tuft)
Tumor cE11 (Enteroendocrine)
cB1 (B IGD+IgM+)
cB2 (B GC-like)
cB3 (B CD40+ GC-like)
cE01 (Stem_TA-like)
cE02 (Stem_TA-like_Immature Goblet)
cE03 (Stem_TA-like prolif)
cE04 (Enterocyte 1)
cE05 (Enterocyte 2)
cE06 (Immature Goblet)
cE07 (Goblet_Enterocyte)
cE08 (Goblet)
cE09 (Best4)
cE10 (Tuft)
cE11 (Enteroendocrine)
cM01 (Monocyte)
cM02 (Macrophage-like)
cM03 (DC1)
cM04 (DC2)
cM05 (DC2 C1Q+)
cM06 (DC IL22RA2)
cM07 (pDC)
cM08 (AS-DC)
cM09 (mregDC)
cM10 (Granulocyte)
cMA01 (Mast)
cP1 (Plasma IgA)
cP2 (Plasma IgG)
cP3 (Plasma IgG prolif)
cS01 (Endo arterial)
cS02 (Endo capillary)
cS03 (Endo capillary)
cS04 (Endo)
cS05 (Endo venous)
cS06 (Endo lymphatic)
cS07 (Endo capillary-like)
cS08 (Endo arterial-like)
cS09 (Endo)
cS10 (Endo tip cells)
cS11 (Endo proif)
cS12 (Endo)
cS13 (Endo venous-like)
cS14 (Endo)
cS15 (Pericyte)
cS16 (Pericyte)
cS17 (Pericyte)
cS18 (Pericyte)
cS19 (Pericyte)
cS20 (Pericyte prolif)
cS21 (Fibro stem cell niche)
cS22 (Fibro stem cell niche)
cS23 (Fibro BMP-producing)
cS24 (Fibro BMP-producing)
cS25 (Fibro CCL8+)
cS26 (Myofibro)
cS27 (CXCL14+ CAF)
cS28 (GREM1+ CAF)
cS29 (MMP3+ CAF)
cS30 (CAF CCL8 Fibro-like)
cS31 (CAF stem niche Fibro-like)
cS32 (Smooth Muscle)
cS33 (Schwann)
cTNI01 (CD4+ IL7R+)
cTNI02 (CD4+ IL7R+SELL+)
cTNI03 (CD4+ IL7R+HSP+)
cTNI04 (CD4+ IL7R+CCL5+)
cTNI05 (CD4+ IL17+)
cTNI06 (CD4+ TFH)
cTNI07 (CD4+ CXCL13+)
cTNI08 (CD4+ Treg)
cTNI09 (CD4+ Treg prolif)
cTNI10 (CD8+ IL7R+)
cTNI11 (CD8+GZMK+)
cTNI12 (CD8+ IL7R+)
cTNI13 (CD8+ T IL17+)
cTNI14 (CD8+ CXCL13+)
cTNI15 (CD8+ CXCL13+ HSP+)
cTNI16 (CD8+ CXCL13+ prolif)
cTNI17 (gd-like T)
cTNI18 (gd-like T PDCD1+)
cTNI19 (gd-like T prolif)
cTNI20 (PLZF+ T)
cTNI21 (PLZF+ T prolif)
cTNI22 (cTNI22)
cTNI23 (NK CD16A+)
cTNI24 (NK GZMK+)
cTNI25 (NK XCL1+)
cTNI26 (ILC3)
PODN
LMOD1
ACTG2
DES
MYLK
SYNPO2
PDLIM3
COL12A1
SVIL
KCNMA1
SORBS1
C11orf96
CKB
SYNM
TGFB1I1
CNN1
FXYD3
HSPB6
PPP1R14A
DSTN
PRDM2
RABGAP1L
VAMP8
EIF2AK3
CXCL13
HIST1H3H
IKZF1
TRBC2
EML3
RBM14
POU2AF1
ZFC3H1
CIITA
CORO1A
SSH2
CD79A
CD37
SLC35C2
SUN2
10
8
6
4
2
0

**Figure S6 (previous page). Heatmap of regressed gene expression in colorectal cancer single-cell reference (Pelka et al, 2021).**

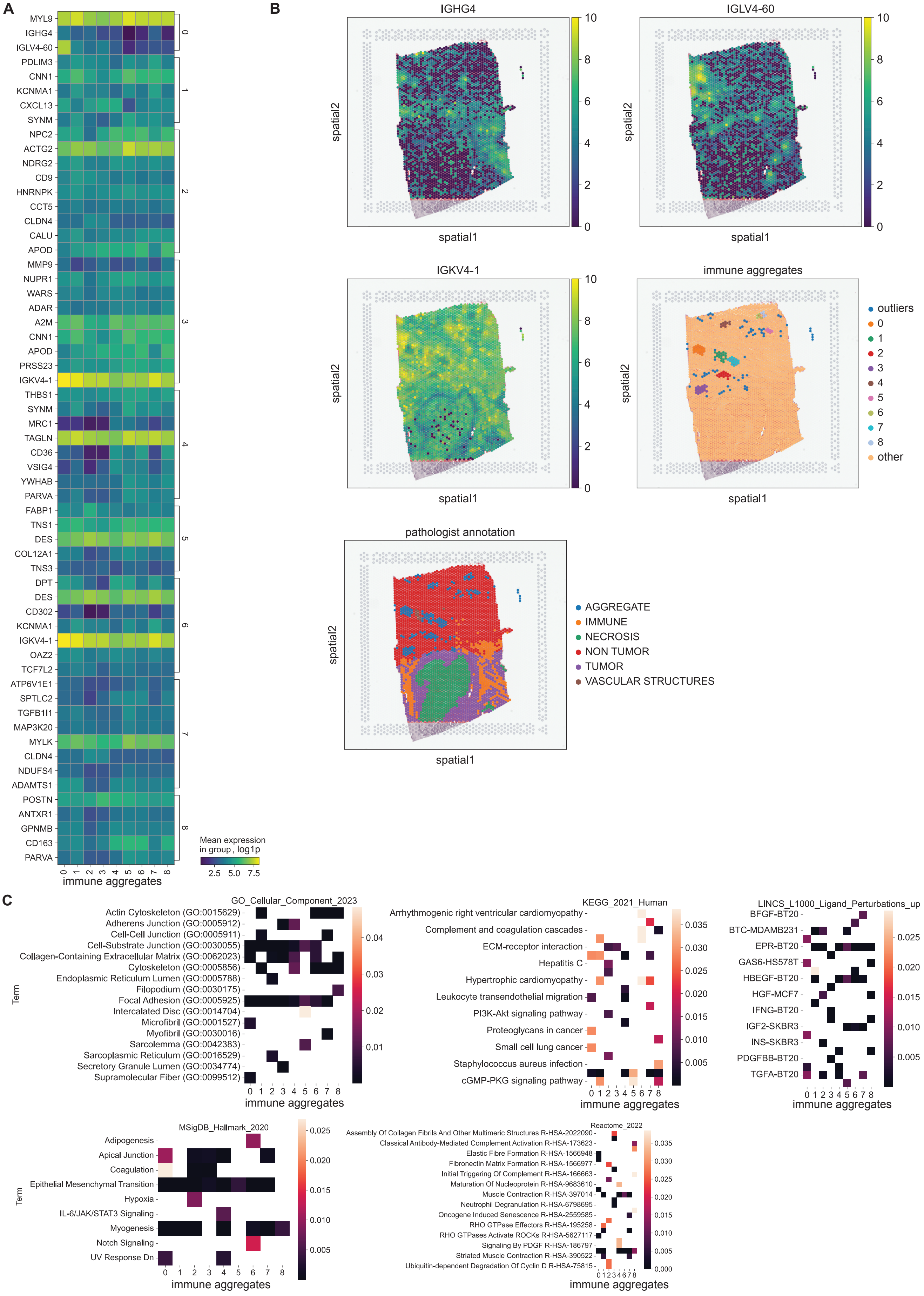
A
MYL9
IGHG4
IGLV4-60
PDLIM3
CNN1
KCNMA1
CXCL13
SYNM
NPC2
ACTG2
NDRG2
CD9
HNRNPK
CCT5
CLDN4
CALU
APOD
MMP9
NUPR1
WARS
ADAR
A2M
CNN1
APOD
PRSS23
IGKV4-1
THBS1
SYNM
MRC1
TAGLN
CD36
VSIG4
YWHAB
PARVA
FABP1
TNS1
DES
COL12A1
TNS3
DPT
DES
CD302
KCNMA1
IGKV4-1
OAZ2
TCF7L2
ATP6V1E1
SPTLC2
TGFB1I1
MAP3K20
MYLK
CLDN4
NDUFS4
ADAMTS1
POSTN
ANTXR1
GPNMB
CD163
PARVA
immune aggregates
Mean expression in group, log1p
2.5 5.0 7.5
B
IGHG4
IGLV4-60
IGKV4-1
immune aggregates
pathologist annotation
spatial1
spatial2
outliers
other
AGGREGATE
IMMUNE
NECROSIS
NON TUMOR
TUMOR
VASCULAR STRUCTURES
C
GO_Cellular_Component_2023
Actin Cytoskeleton (GO:0015629)
Adherens Junction (GO:0005912)
Cell-Cell Junction (GO:0005911)
Cell-Substrate Junction (GO:0030055)
Collagen-Containing Extracellular Matrix (GO:0062023)
Cytoskeleton (GO:0005856)
Endoplasmic Reticulum Lumen (GO:0005788)
Filopodium (GO:0030175)
Focal Adhesion (GO:0005925)
Intercalated Disc (GO:0014704)
Microfibril (GO:0001527)
Myofibril (GO:0030016)
Sarcolemma (GO:0042383)
Sarcoplasmic Reticulum (GO:0016529)
Secretory Granule Lumen (GO:0034774)
Supramolecular Fiber (GO:0099512)
Term
immune aggregates
KEGG_2021_Human
Arrhythmogenic right ventricular cardiomyopathy
Complement and coagulation cascades
ECM-receptor interaction
Hepatitis C
Hypertrophic cardiomyopathy
Leukocyte transendothelial migration
PI3K-Akt signaling pathway
Proteoglycans in cancer
Small cell lung cancer
Staphylococcus aureus infection
cGMP-PKG signaling pathway
LINCS_L1000_Ligand_Perturbations_up
BFGF-BT20
BTC-MDAMB231
EPR-BT20
GAS6-HS578T
HBEGF-BT20
HGF-MCF7
IFNG-BT20
IGF2-SKBR3
INS-SKBR3
PDGFBB-BT20
TGFA-BT20
MSigDB_Hallmark_2020
Adipogenesis
Apical Junction
Coagulation
Epithelial Mesenchymal Transition
Hypoxia
IL-6/JAK/STAT3 Signaling
Myogenesis
Notch Signaling
UV Response Dn
Reactome_2022
Assembly Of Collagen Fibrils And Other Multimeric Structures R-HSA-2022090
Classical Antibody-Mediated Complement Activation R-HSA-173623
Elastic Fibre Formation R-HSA-1566948
Fibronectin Matrix Formation R-HSA-1566977
Initial Triggering Of Complement R-HSA-166663
Maturation Of Nucleoprotein R-HSA-9683610
Muscle Contraction R-HSA-397014
Neutrophil Degranulation R-HSA-6798695
Oncogene Induced Senescence R-HSA-2559585
RHO GTPase Effectors R-HSA-195258
RHO GTPases Activate ROCKs R-HSA-5627117
Signaling By PDGF R-HSA-186797
Striated Muscle Contraction R-HSA-390522
Ubiquitin-dependent Degradation Of Cyclin D R-HSA-75815

**Figure S7 (previous page). Regression analysis around individual immune aggregates unveils antibody gene enrichment outside the immune aggregate compartment.**

(A) Top genes enriched in the neighborhood of individual immune aggregates. Only genes passed Mann-Whitney U test 'one vs other' with Benjamini-Hochberg multiple correction are shown (p-value<0.05). (B) Top two rows show three found antibody genes and their expression pattern across the histological slide. Second and third panels show the annotation of individual immune aggregates and the pathologist' annotation of the slide. (C) Results of pathway analysis across all individual immune aggregates. Top 50 genes were passed to the gene set enrichment analysis.
